# Active antennal movements in *Drosophila* tune wind encoding

**DOI:** 10.1101/2022.09.30.510392

**Authors:** Marie P Suver, Ashley M Medina, Katherine I Nagel

## Abstract

Many insects actively move their antennae, but how these movements influence sensory encoding is not fully understood. Antennae are used to smell odors^1,2^, detect auditory cues^3,4^, and sense mechanosensory stimuli such as wind^5^ and objects^6–8^, frequently by combining active movement with sensation. Genetic access to antennal motor systems would thus provide a powerful tool for dissecting the circuit mechanisms underlying active sensing, but little is known about how the most genetically tractable insect, *Drosophila melanogaster*, moves its antennae. Here we use DeepLabCut to measure how tethered *Drosophila* move their antennae in the presence of sensory stimuli, and identify genetic reagents for controlling antennal movement. We find that flies perform both slow and fast antennal movements in response to wind-induced deflections, but not the attractive odor apple cider vinegar. We describe four muscles in the first antennal segment that control antennal movements, and identify genetic driver lines that provide access to two groups of antennal motor neurons and an antennal muscle. Through optogenetic inactivation, we provide evidence that antennal motor neurons are specialized for different movement speeds. Finally, we show that activation of antennal motor neurons and muscles can improve the gain and acuity of wind direction encoding. Together, our experiments provide insight into the neural control of antennal movement and suggest that *Drosophila* actively position their antennae to tune the precision of wind encoding.

## Results

### Flies exhibit diverse active movements that are promoted by wind and flight

To measure active antennal movements in *Drosophila*, we presented head-fixed flies with wind stimuli of various speeds (0-200 cm/s) and directions (−90°, -45°, 0°, +45°, and +90°) while recording antennal movement using a camera positioned in front of the fly (Fig. 1A). We used DeepLabCut^9^ to track the position of multiple points on the head and antennae (Fig. S1A). We then used a subset of these points to compute the fly’s midline and the angles of the second and third antennal segments in 2 dimensions (Fig. 1B). We observed both passive deflections of the third antennal segment that were tightly locked to the wind stimulus (Fig. 1C, left), and active movements that appeared as simultaneous deflections in both the second and third segment (Fig. 1C, right). Because antennal muscles control the joint between the first and second segments^10^, we interpret these second segment movements as ‘active’ movements controlled by the fly’s motor system. As observed in previous studies^11–13^, third segment (passive) movements were rapid, nearly tonic, and depended systematically on wind speed and direction (Fig. 1D,E, top). These movements are detected by mechanosensitive Johnston’s organ neurons (JONs^14–16^), and enable walking flies to determine wind speed and direction^11,12^. In contrast, second segment (active) movements were more variable across trials and flies (Fig. 1D,E, bottom).

**Figure 1.**
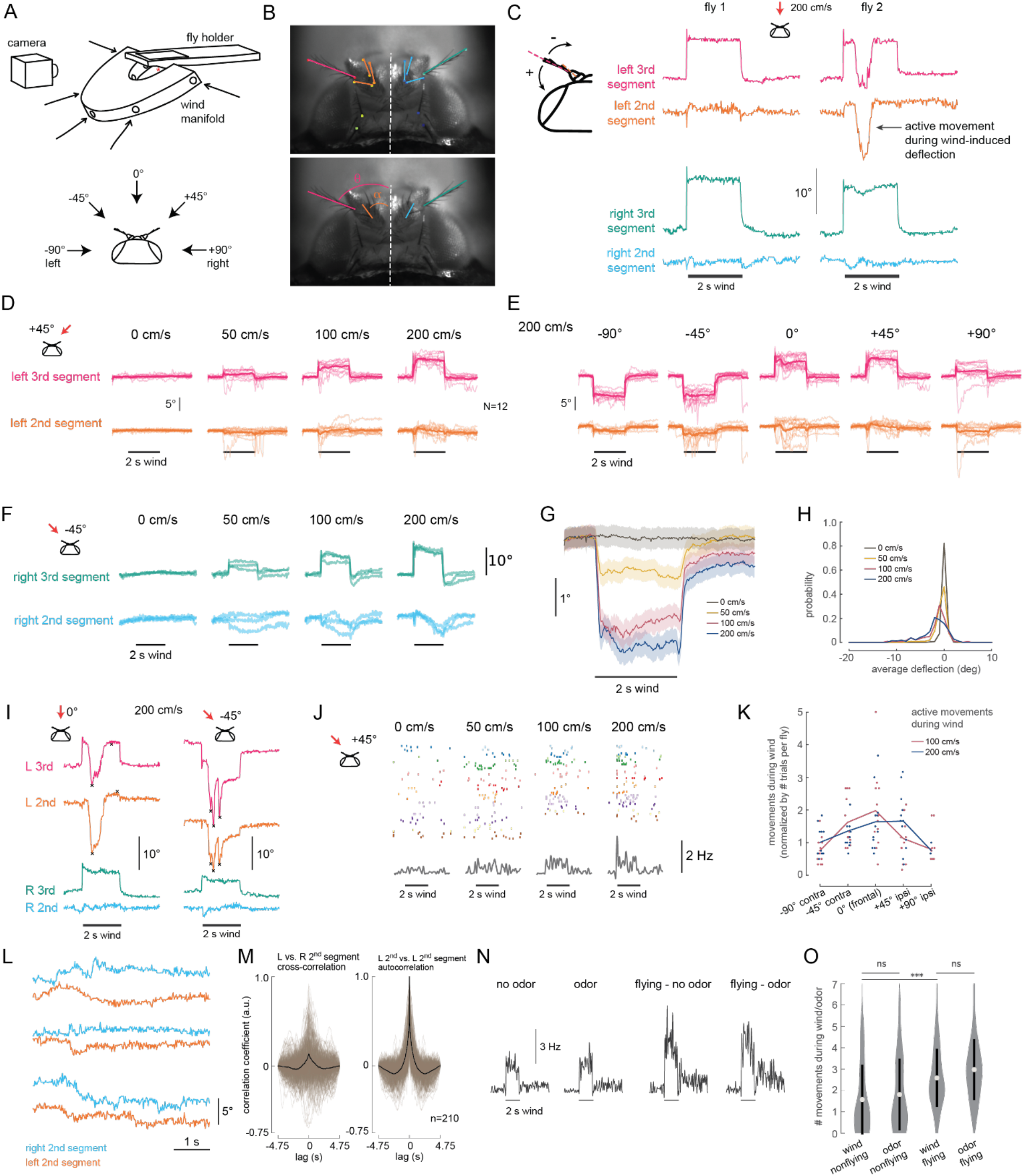
Active antennal motions in response to wind stimuli. (A) Schematic of experimental setup showing wind directions and camera position. (B) Still images of the fly head showing points (top) and angles (bottom) tracked. The base of two pairs of cephalic hairs were used to determine the midline axis of the head (dashed white line). Three hairs (base and tip) on the second antennal segment were used to determine the angles of the second segments (orange and blue lines). The arista base and tip were used to determine the angles of third segments (pink and green lines). (C) Two single-trial examples of antennal angles for all measured segments are shown. Colors as in (B). A 2 s frontal (0°) wind stimulus elicits posterior (positive) deflections of both third antennal segments. Left traces: Passive movement deflects the third segments only. Right traces: an active movement of the left antennae is visible in both third and second segments on one side. Schematic shows antennal deflection sign convention in which positive deflections indicate posterior deflections of the antenna, and negative indicate anterior deflections. (D) Average antennal deflections for the left second and third segments in response to a 2 s contralateral wind (+45°) stimulus across windspeeds. Single fly averages (thinner lines) and cross-fly averages (thicker lines) are plotted. (E) Average antennal deflections for the left second and third segments in response to 200 cm/s wind across five directions. Responses in (D) and (E) are plotted on the same scale for the same set of 12 flies. (F) Single-trial examples from one fly exhibiting slow adaptive movements in the second segment. Magnitude of the adaptive movement increases with windspeed. (G) Time course of average second segment deflection across windspeeds. Error bars show standard error of the mean across 420 trials from 12 flies. (H) Distribution of steady-state second segment deflections for each windspeed. Larger deflections are observed at higher windspeeds. (I) Examples of rapid active movements in one fly. Detected active movements are marked (see Methods and Fig. S1B, C). (J) Raster of active antennal movements (both antennae) for different speeds of a single wind direction (45°). Different colors represent trials from different flies (N = 12 non-flying flies). Peristimulus time histogram of active movements shown in gray below raster. (K) Average number of active movements per trial as a function of wind direction. Dots represent single flies (N= 12), lines represent average across flies. (L) Single trial examples of correlated movements in the left (orange) and right (blue) second segments in the absence of wind. (M) Cross-correlation of left and right second segments for n = 210 no-wind (0 cm/s) trials from N=12 flies and auto-correlation of left segments only. Single trial correlations in gray, cross-trial average in black. Half maximum width of the cross-correlogram is 0.77+/-1.3 s, and of the auto-correlogram is 0.20+/-0.14 s. Auto-correlogram is significantly narrower (p=0.011, paired student’s t-test). (N) Frequency of active movements in odorized (1% apple cider vinegar) versus non-odorized wind, in both non-flying and flying flies. (O) Violin plots showing the average number of active movements during the wind stimulus (circles), the standard deviation (black line), and the range (shaded region). For each condition shown, average number of movements per second during the stimulus was 1.57+/3.20, 1.81+/-2.50, 2.59+/-3.96, and 2.98+/-4.40, respectively. (N) Flying flies respond with more active movements regardless of the presence of odor (p<0.0001, student’s t-test), whereas no significant change in movements is observed in the presence of odor (p=0.15 and p-0.072 during nonflying and flying trials, respectively). Flight data is from N = 6 of the 7 total flies in (L) and (M). Averages represent the mean of n = 206, 198, 82, and 88 trials, respectively. All flies in Figure 1 are Canton-S > Chrimson and experiments were performed in the presence of red light to serve as controls for later optogenetic activation experiments.

We observed two main kinds of active antennal movements: slow, adaptive movements (Fig. 1F-H), and more rapid flick-like movements (Fig. 1I-K), both of which were promoted by wind. Slow movements typically occurred in response to backward/posterior deflection of the third segment (increased third segment angle), leading to a forward movement of both segments over time (decreased second and third segment angles over time, example in 1F). Across all wind stimuli, we observed forward deflections (decreased angle) of the second segment on average, with larger movements in response to faster windspeeds (Figure 1G-H). Rapid movements (examples in Fig. 1I) were also preferentially triggered during or after a wind stimulus, with the frequency of movements increasing with wind speed (Fig. 1J) and frontal direction (Fig. 1K, Supplemental Video 1). Taken together, these data suggest that wind promotes active movements through large deflections of the third segment.

In our experiments, active antennal motions were sometimes much larger in one antenna than the other (e.g. Fig. 1C). However, they were also occasionally coordinated between the left and right antennae. To examine the correlation between left and right antennal movements, we plotted the cross-correlogram of second segment movements during the no-wind condition (Fig. 1L, M, S1D). The cross-correlogram has a small peak centered at 0, indicating that the two antennae can move together. This peak is slightly wider, but lower amplitude, than the auto-correlogram of second segment movement (Fig. 1M), suggesting that fast movements have a characteristic timescale, with a weak correlation between the two antennae.

In several arthropods, such as locusts^1^ and cockroaches^17^, antennal movements increase in the presence of odor and are thought to function like sniffing^18^ to increase the persistence of odorants at olfactory sensilla. To determine if *Drosophila* alter olfaction through active antennal movements, we compared antennal movements in response to odorized and non-odorized wind. Surprisingly, we found no difference between these two conditions in *Drosophila* (Fig. 1N). In contrast, we did observe an increase in active movements when flies began to fly (average increase of 1.013 movements/s, p<0.0001, student’s t-test; Fig. 1O, Supplemental Video 2) as previously observed^19^. Further, blocking active movements by stabilizing the first and second segments of the antennae did not impair odor-evoked upwind walking (Fig. S1E). Together, these data argue that active antennal movements in *Drosophila* do not function to improve odor detection and are not required for wind orientation during walking, at least in the presence of laminar airflow. Rather, they suggest a possible role in detecting dynamic air currents in flight, when effective airspeeds are higher and antennal deflections are larger.

### Distinct motor neurons promote antennal movements with different time courses

While mechanosensory neurons of the fly antenna have been extensively characterized^14–16^, little is known about the muscles and motor neurons controlling active antennal movements in *Drosophila*. To investigate motor control of the antennae, we began by visualizing antennal muscles and tendons (see Methods, Fig. 2A-C). Although a classical study suggested that there are only two muscles in the first antennal segment^10^, we observed four distinct muscles within this structure (Fig. 2B, C). Tendon labeling (Fig. 2C) indicates that the insertion point of these antennal muscles is likely at the interior portion of the second antennal segment, where it joins with the first, thus enabling the fly to move the second segment relative to the first segment.

**Figure 2.**
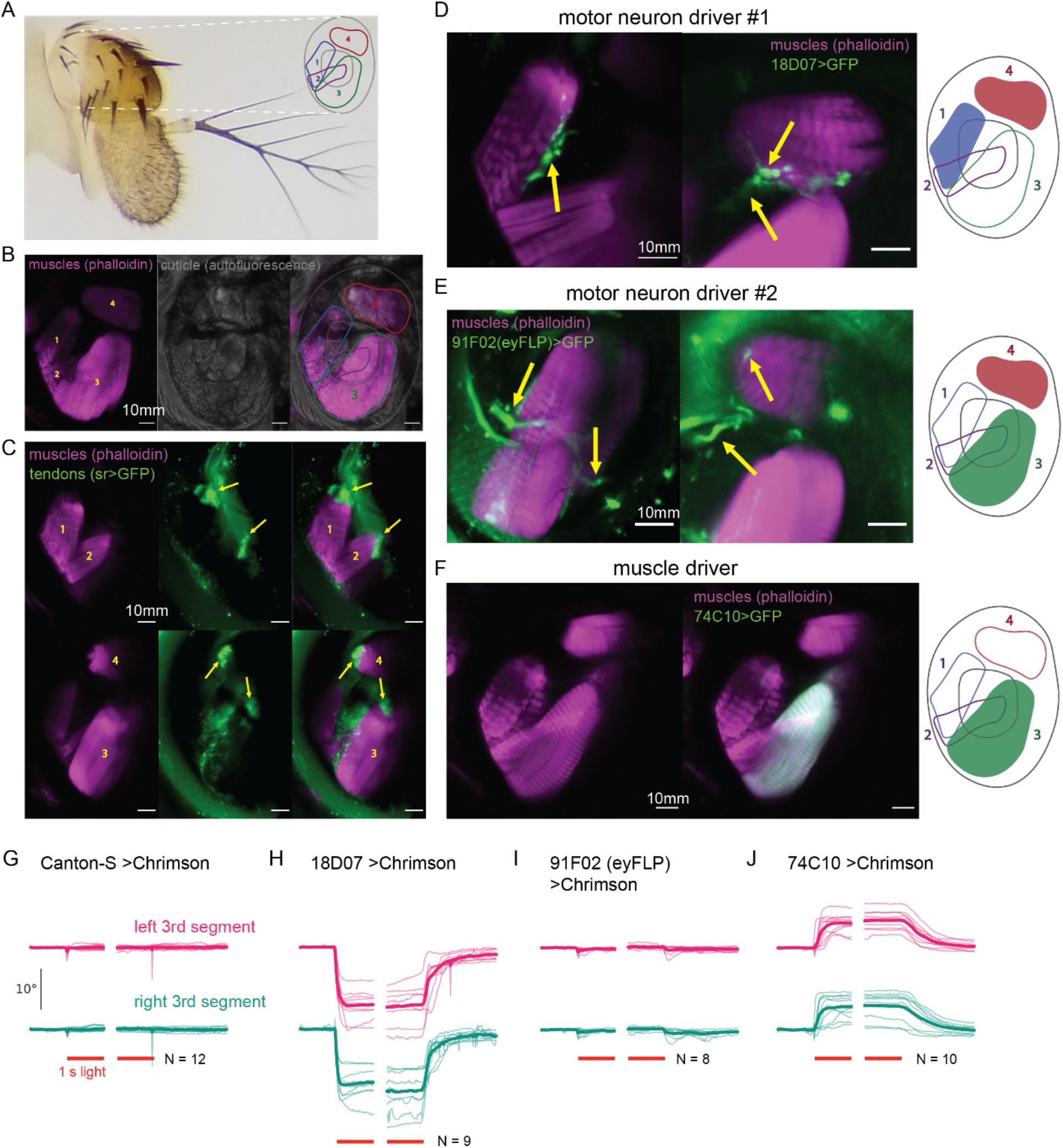
Anatomy of antennal muscles and genetic lines labeling antennal motor neurons. (A) Antennal preparation with first antennal segment outlined as an inset, indicating the location of the four antennal muscles, numbered 1 to 4 from anterior to posterior. Schematic depicts coronal (frontal) projection of the antennae. Gray lines indicate outer and inner cuticle surrounding the first antennal segment. (B) Four muscles in the first antennal segment. Left: phalloidin Alexa 568 stain for muscle (magenta). Center: cuticle (gray). Right: overlay of phalloidin stain and cuticle with outlines of muscles. (C) Tendon expression using *sr-GAL4>10xUAS-GFP* (green) with the four muscles in the first antennal segment. Yellow arrows indicate tendons; muscles numbered as in (A). (D) Motor neuron processes labeled by *18D07*. Left: muscle 1 innervation. Center: muscle 4 innervation. Right: schematic of innervated muscles. (E) Motor neuron processes labeled by *91F02*. Left: muscle 3 innervation. Center: muscle 4 innervation. Right: schematic of innervated muscles. (F) Muscle labeled by *74C10*. Left: phalloidin stain of muscles. Center: GFP expression overlaid on phalloidin stain. Right: schematic of labeled muscle. (G-J) Antennal deflections produced by Chrimson activation in control flies (Canton-S>Chrimson, N=12 flies) and three experimental lines (*18D07*, N=9 flies; *91F02*, N=8 flies; and *74C10*, N=10 flies). Thin traces represent average response of one fly, thick traces are means across flies. Anatomy and activation for *91F02* only used eyFLP to suppress expression in JONs in this line (see Methods).

Next, we used the FlyLight collection^20^ to search for genetic lines that label antennal motor neurons, selecting candidate lines with expression in the antennal mechanosensory and motor center (AMMC) where motor neuron input processes reside^21,22^, and in the lateral margin of the antennal nerve, where antennal motor neuron axons run^23–25^. We then screened candidate lines for anatomical innervation of antennal muscles (Fig. 2D-F) and for antennal deflections during optogenetic activation (Fig. 2G-J, S2A-D). Through these approaches, we identified two lines labeling antennal motor neurons (Fig. 2D, E, H, I) and one line labeling an antennal muscle (Fig. 2F, J).

One motor neuron driver (*18D07-GAL4*, Fig. 2D) labeled two motor neurons synapsing onto the dorso-medial antennal muscles 1 and 4. Consistent with this anatomy, activation of this line drove movement of the antennae forward and up (decreased antennal angle, Fig. 2H, Supplemental Video 4). A second motor neuron driver (*91F02-GAL4*, Figure 2E) labeled motor neurons innervating lateral muscles 3 and 4. Curiously, activation of this line evoked only transient antennal movements at light onset and offset (Fig. 2I, Supplemental Video 5). This line also labeled a subset of JONs; we used *eyFLP>tub(FRT*.*stop)Gal80* to suppress expression in these neurons for anatomical and activation experiments^26^. Finally, one driver produced large antennal deflections down and backward (positive antennal angle, Fig. 2J), but did not label any motor neurons. Instead, this driver (*74C10-GAL4*, Fig. 2F, Supplemental Video 6) labeled the large ventro-lateral muscle 3 which pulls the antennae down and back. Thus, these three lines allow for differential experimental control over antennal positioning.

To determine the role of these motor neurons in active movements, we expressed the inhibitory opsin GtACR*1*^27^ in each line, and measured wind-evoked second segment (active) movements in the presence and absence of light. In control flies (*empty-GAL4>GtACR1*), light did not alter the average time course or frequency of active movements (Fig. 3A, S3A, E, F, Supplemental Video 7), although it slightly reduced mean second segment deflection (Fig. S3E). For this reason, we subsequently compared active movements in motor-neuron-silenced flies to those of control *empty-GAL4>GtACR1* flies in the presence of light.

**Figure 3.**
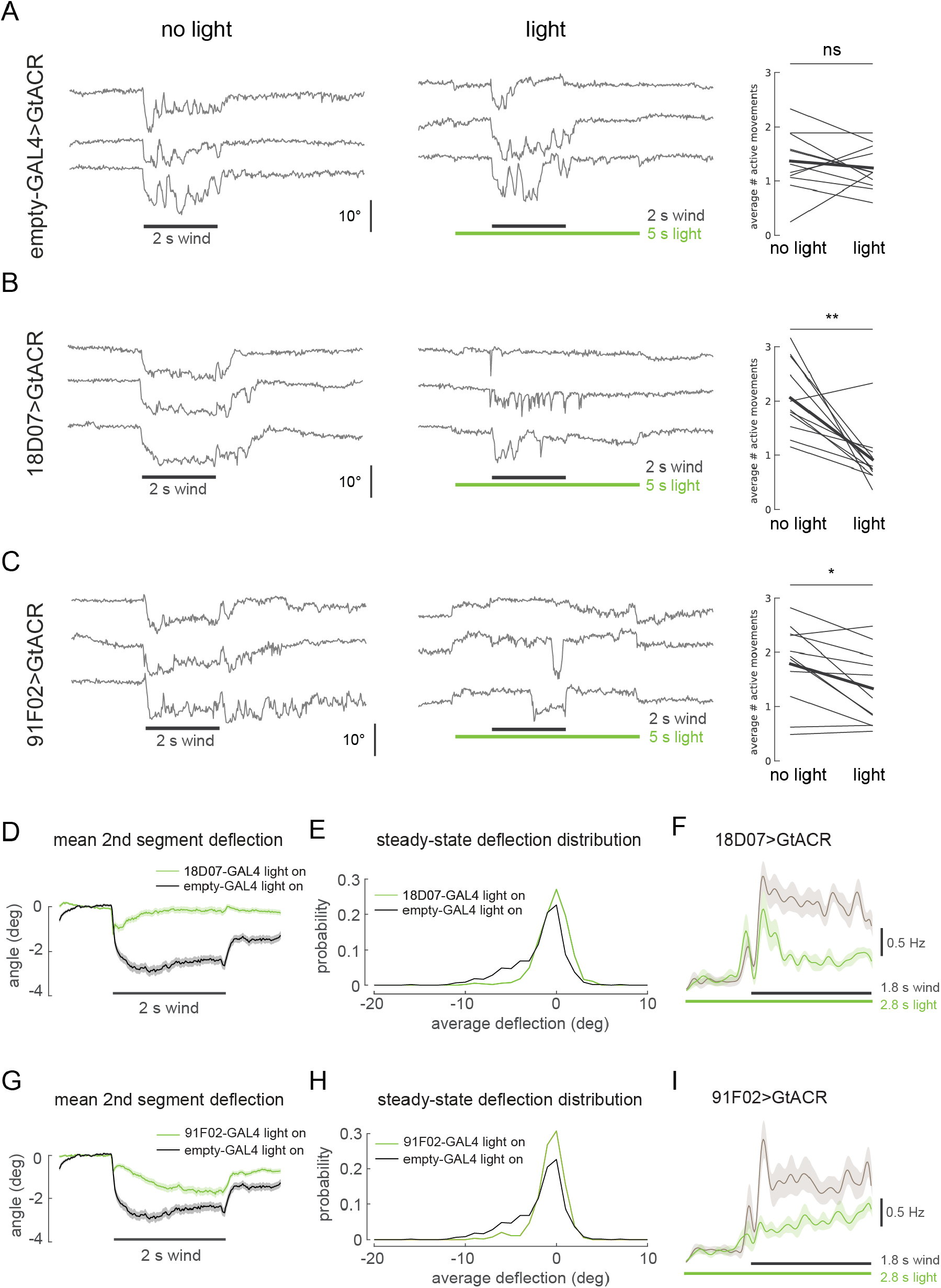
Motor neurons contribute to active movements on different timescales. (A) Effects of optogenetic silencing on active movements in control flies (*empty>GtACR1*). Left: example second segment traces in response to wind (45° contralateral, 200cm/s) with no light. Center: example 2^nd^ segment traces in response to wind with light. Right: No significant difference in the number of active movements in light vs no light (average difference = -0.128+/-0.503 std, p=0.441, N=10 flies). Thin lines represent single fly averages; thick line represents cross-fly average. (B) Effects of optogenetic silencing of *18D07* on active movements. Left: examples, wind with no light. Center: examples, wind with light. Right: Significant decrease in active movement number with *18D07* silencing (average difference = -1.138+/-0.871, p=0.002, N= 11 flies). (C) Effects of optogenetic silencing of *91F02* on active movements. Left: examples, wind with no light. Center: examples, wind with light. Right: Significant decrease in active movement number with 91F02 silencing (average difference = -0.454+/-0.482, p=0.014, N=11 flies). All p-values for (A-C) from paired t-test. (D) Time course of average 2^nd^ segment deflection in *18D07*-silenced flies versus control-silenced flies (both in the presence of light). (E) Distribution of steady-state antennal deflections in the same flies. (F) Peri-stimulus time histogram of active antennal movements in the same flies. (G-I) Same as D-F for *91F02* silencing. Error bars in (D-I) represent standard error of the mean across trials.

Silencing the first motor neuron line (*18D07>GtACR1*) significantly reduced the frequency of active movements, and altered their time course (Fig. 3B, S3 B, G, H, Supplemental Video 8). In the absence of light, active movements often persisted for the duration of the wind stimulus and beyond. During light, the remaining active movements were rapid and clustered near the beginning of the wind stimulus. To quantify these observations, we examined the mean second segment deflection across wind directions (Fig. 3D, E) and computed the peristimulus time histogram of active movement times (Fig. 3F). Silencing *18D07* motor neurons abolished the tonic deflection observed in control flies (Fig. 3D, E), and most of the remaining active movements were clustered near the beginning of the wind stimulus (Fig. 3F). Silencing of *18D07* motor neurons also produced a tonic offset in second segment position (Fig. S3B, D), suggesting that these neurons contribute to steady-state antennal positioning. Together, these data suggest that *18D07* motor neurons contribute to slow active movements and tonic antennal positioning.

Silencing the second motor neuron line (*91F02>GtACR1*) also reduced active movement frequency. However, the remaining active movements were longer in duration and could occur at any point during the stimulus (Fig. 3C,S3C, I, J, Supplemental Video 9). Both the mean deflection (Fig. 3G,H) and the peri-stimulus time histogram (Fig. 3I) showed a ramping profile, indicating that the remaining movements preferentially occurred later in the stimulus than in *18D07* silencing. Silencing *91F02* also produced tonic changes in antennal position, of a similar magnitude to those produced by *18D07* silencing (Fig. S3C, D). It remains possible that some of these effects are due to silencing of JONs in this line, however, these results suggest that *91F02* contributes more to rapid movements, particularly those triggered by wind onset.

### Active antennal movements alter the gain and precision of wind encoding

Flies and other walking insects compute wind direction from differential displacements of their antennae, which are decoded by central neurons in the brain to subserve wind orientation behavior^5,11–13,28–30^. Because this differential displacement depends on antennal angle, we reasoned that changing antennal position should change peripheral encoding of wind direction. To test this hypothesis, we expressed Chrimson in each of our motor neuron and muscle lines, and compared the encoding of wind direction by antennal displacement differences to a genetic control (*Canton-S>Chrimson*, Fig. 4A-D).

**Figure 4.**
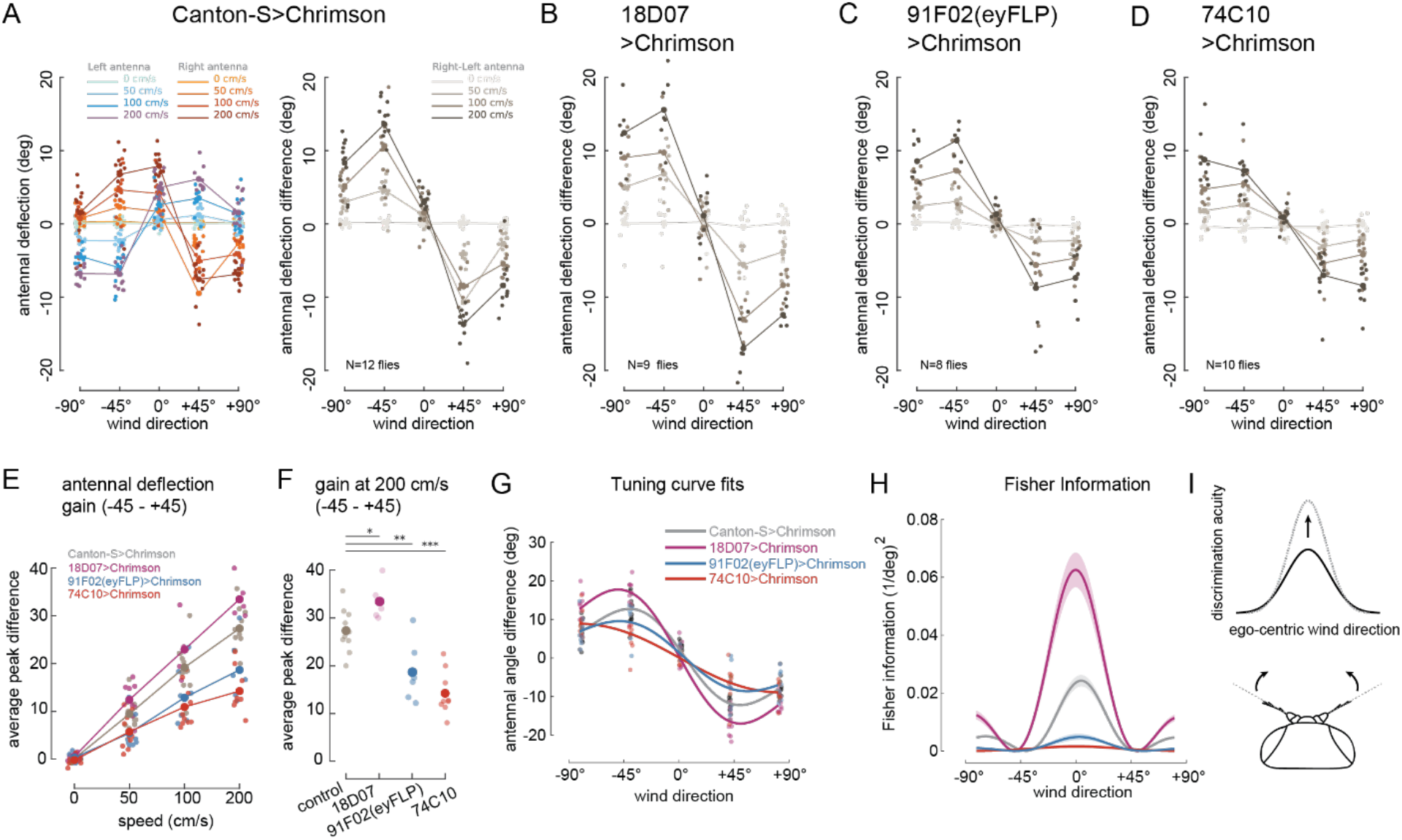
Antennal positioning modifies the gain and precision of wind encoding. (A) Steady-state third segment (passive) antennal deflection across wind directions and speeds in control flies (*Canton-S>Chrimson*). Left: blue-purple hues indicate left antennal responses, orange-red hues indicate right antennal responses. Right: difference between right and left antennal deflections shown in gray hues. Dots represent single flies (N=12 flies). Lines represent cross-fly averages. (B) Effects of activating *18D07* on wind encoding by third segment displacement differences (N=9 flies). (C) Effects of activating *91F02* on wind encoding by third segment displacement differences (N=8 flies). For these experiments, we suppressed expression in JONs using eyFLP (*eyFLP>GAL80;91F02>Chrimson*, see Methods). (D) Effects of activating *74C10* on wind encoding by third segment displacement differences (N=10 flies). Colors and symbols in (B-D) as in (A). (E) Gain (average 3^rd^ segment displacement difference at -45° minus +45°) during light activation across speeds for the four genotypes. (F) Antennal deflection gain (average at -45° minus +45°) across flies during light activation at 200 cm/s (subset of data in (E)). Gain significantly changes with *18D07* activation (p=0.025), *91F02* activation (p=0.007), and *74C10* activation (p<0.0001). Comparison made with two-sided student’s t-test. (G) Tuning curves (see Methods) fit to third segment displacement differences for each genotype at 200 cm/s. (H) Fisher information computation for each genotype at 200 cm/s. Shaded region indicates standard error of the mean. (I) Hypothesized function of active antennal movements. Acuity of wind direction encoding relative to the fly’s body axis increases by actively positioning the antennae.

As described above, activation of the *18D07* motor neuron line moved the antennae forward and up, similar to natural movements induced by the start of flight (Supplementary Video 2). Wind-evoked displacements of the third segment were typically larger in this position, leading to a wind direction tuning curve with higher gain (Fig. 4B). In contrast, while activation of the *91F02* motor neuron line produced little net change in antennal position (Fig. 2I), 91F02 activation reduced the gain of wind encoding slightly (Fig. 4C). Finally, activation of the muscle driver line *74C10* pulled the antennae down and away from the midline, roughly the opposite direction from flight position. This manipulation reduced wind encoding gain and shifted the tuning curve so that the peak difference between antennal displacements occurred at more eccentric wind angles (+/-90° instead of +/-45°, Fig. 4D). Taken together, these results support our hypothesis that active changes in antennal position alter the encoding of wind direction at the periphery (Fig. 4E, F).

Do these changes in gain and tuning alter the information the fly can encode about wind direction? To address this question, we computed the Fisher Information, a measure of discriminability between nearby stimulus directions^30^ during activation for each genetic line. The Fisher Information is equal to the slope of the tuning curve normalized to the standard deviation of the response, and measures how much the response changes (relative to its variability) when the stimulus changes. To compute the Fisher Information, we fit a smooth function to the displacement tuning curves for the fastest wind direction (Fig. 4G), differentiated this function, and divided by the standard deviation across trials for each genotype (see Methods). This analysis (Fig. 4H) shows that *18D07* activation strongly increases the Fisher Information for frontal wind directions, whereas *91F02* and muscle activation (*74C10*) decrease Fisher Information. These effects arise both because of the changes in tuning curve slope, and because responses during *18D07* activation were more reliable across trials. Flies can thus use active antennal positioning to tune their acuity for wind direction (Fig. 4I).

## Discussion

Many animals actively move their sensors to alter the information they can obtain from their environment. For example, humans actively move their hands and fingers to determine the size and texture of objects^31,32^, rodents whisk to learn about their surroundings in the dark^33^, and both mammals and lobsters sniff to alter the flow of olfactory information^18,34^. While such active movements are generally proposed to increase sensory information or acuity, this can be challenging to measure quantitatively, as it requires experimental control over animal movement.

Insect antennal movements provide an attractive model for studying active sensing. Many insects move their antennae to explore objects in their environment ^7,8^, in response to attractive odors^1^, or during the transition to flight^21,35^. Antennal movements reflect input from the olfactory^17^, mechanosensory^6^, and visual systems^36,37^, providing an excellent model for studying multisensory integration. A variable number of antennal muscles have been described in different insects, and each is innervated by motor neurons originating in the antennal mechanosensory and motor center ^22,23,24^ (AMMC^24,25,38^). These motor neurons receive fairly direct synaptic input from mechanosensory neurons at each of the antennal joints (JONs and Böhm’s bristles ^21,22^, as well as indirect input from olfactory^17^, visual^36,37^, and descending motor systems^39^. While many studies have investigated multisensory control of antennal positioning in insects, no studies have yet provided genetic access to antennal motor circuitry.

In this study, we set out to characterize antennal movements in the genetic model organism *Drosophila melanogaster*, and to identify genetic lines that would provide experimental control over antennal movement. Using machine learning, we characterized antennal movements in response to wind and odor stimuli. Surprisingly, we found that fruit flies do not increase antennal movements in response to an attractive odor. Instead, active movements seem to be driven most strongly by wind and during flight. We observed slow adaptive movements and more rapid flicking movements, both evoked by large antennal deflections, but often outlasting this mechanical stimulus. We also confirmed previous observations (also seen in other insects) that flies move their antennae forward toward the midline during flight^21,35^. These observations suggest that flies may use antennal movement primarily to tune mechanosensory input, such as wind sensing, rather than to tune olfactory input, as has been proposed in other arthropods^1,18^.

In support of this hypothesis, we showed that experimentally altering antennal position, by activating either antennal motor neurons or antennal muscles, could alter the gain and shape of the antennal wind encoding tuning curve. Moving the antennae forward toward the midline, as insects do during flight, increases frontal wind tuning gain and acuity. This may represent an adaptation to the higher airspeeds a fly encounters during flight (typically 0.5-1 m/s^40^) in contrast to walking, where windspeeds above ∼15-20 cm/s cause freezing behavior^13^. During flight, airflow primarily originates from frontal directions, as any external wind vector will be summed with the airspeed produced by the fly’s own forward movement. In contrast, the more lateral position of the antennae during walking may promote sensation of lateral and posterior mechanosensory stimuli, such as auditory signals from conspecifics. Further, although our optogenetic experiments led to symmetric movements of the antennae, and thus “foveation” in a region directly in front of the fly, our measurements of spontaneous antennal movements suggest that the two antennae can be positioned independently. Thus, flies may be able increase wind acuity in different regions of space by differentially positioning their antennae. The function of the rapid flicking movements we observed in response to wind is less clear, although these could act to calibrate wind sensing. Both slow and fast movements decreased after motor neuron silencing, indicating that they represent active movements. However, we cannot rule out the possibility that rapid movements reflect active amplification of the more turbulent airflows experienced during high wind speeds and in flight.

We identified two genetic lines that label 4 motor neurons, innervating 3 out of 4 antennal muscles. Because we did not identify any motor neurons innervating muscle 2, and because antennal muscles in various other insects typically are innervated by more than one motor neuron^24,25,38^, we expect that *Drosophila melanogaster* possesses additional antennal motor neurons. We attempted to determine the total number of antennal motor neurons using a line labeling all motor neurons (*vGlut-GAL4*), but were unable to visualize individual axons in the antenna using this line. Intriguingly, our silencing data suggest that different antennal motor neurons may subserve slow positioning and fast flicking movements. This is similar to observations of fly leg motor neurons^41^ which are organized by movement speed. Examples of organization of motor neurons by movement speed are known across the animal kingdom in species including insects^42,43^, fish^44^, and mammals^45,46^. Future studies, using electron microscopy and x-ray images of the antenna and ventral brain, will allow for a more complete mapping of antennal motor circuitry, and a more detailed understanding of the mechanics of antennal joints.

Although we show here that active antennal positioning enables the fly to dynamically tune its peripheral encoding of wind direction, this presents a challenge for central neurons that decode wind direction to drive orientation behavior^11,12,29^. To compute the true wind direction in the presence of active positioning movements, central neurons might receive a predictive motor signal that would allow them to distinguish passive deflections (which displace the third segment only) from active movements (which displace both third and second segments). In several mechanosensory systems, signaling by neurons downstream of the sensory periphery is suppressed by the motor system^47–49^. Genetic access to central neurons that decode wind direction in *Drosophila* will allow us to pinpoint how this computation is performed at a detailed synaptic level.

## Acknowledgements

We would like to thank Peter Polidoro and Janelia Research Campus for generously providing the tachometer for recording wingbeat signals, Jonathan Victor for advice with the Fisher Information calculation, and Mackenzie Mathis for advice with DeepLabCut. Jonathan Victor, Tony Azevedo, and Floris van Breugel provided helpful feedback on the manuscript. This work was supported by the NIH through R01DC017979 to K.N., by a BRAIN Initiative K99 K99NS114179 to M.P.S., and by a Leon Levy Foundation Fellowship to M.P.S. A.M.M. was supported by a Diversity Supplement to R01DC017979 and by a McKnight Pecot Fellowship.

## Author contributions

M.P.S., A.M.M., and K.I.N designed experiments. A.M.M. and M.P.S. performed anatomy experiments and confocal imaging. M.P.S. and A.M.M. performed behavior experiments. M.P.S. and K.I.N. analyzed data. M.P.S., K.I.N, and A.M.M. wrote the manuscript.

## Inclusion and Diversity Statement

One or more of the authors of this paper self-identifies as an underrepresented ethnic minority in science.

## Figures and figure legends

**Supplementary Figure 1 (related to Figure 1):**
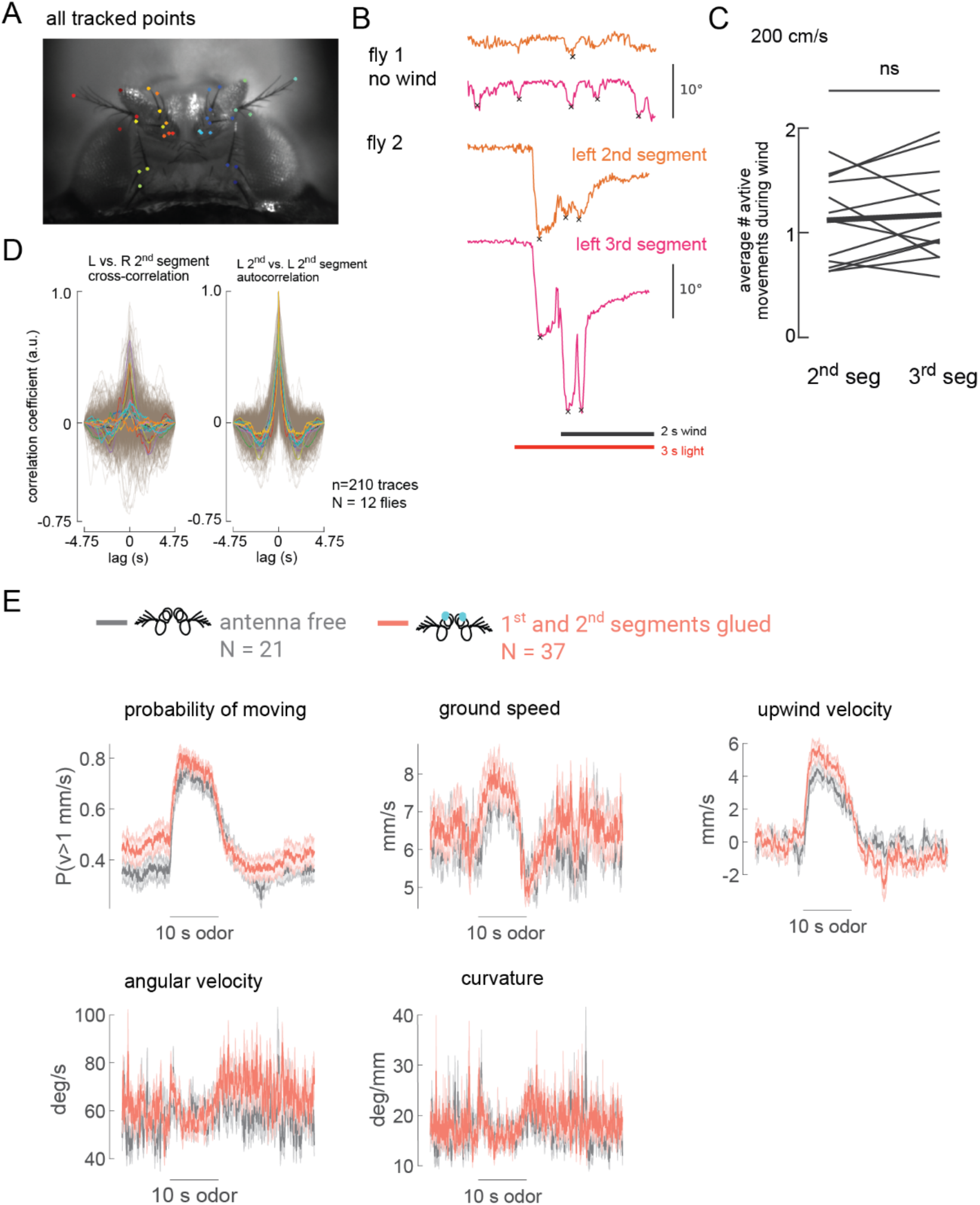
(A) Left: All parts on the head of the fly tracked by the trained network. (B) We used third segment traces to detect rapid active movements (see Methods) as their signal-to-noise was typically higher than second segments. Panel shows example traces with active movements marked, detected using the left second segment (orange) or third segment (pink). (C) Quantification of active movements using the second versus the third antennal segment movements in control flies at 200 cm/s (all directions). NO significant difference between average number of movements detected in the third versus second segments (average difference is 0.051 +/-0.252, p=0.64, paired t-test). (D) Cross-correlation of left and right second segments for n = 210 no-wind (0 cm/s) trials from N=12 flies and autocorrelation of left segments only, with single-fly averages plotted in color. Single trial correlations shown in gray, cross-trial average in black. (E) In freely walking flies, blocking active antennal movements with a small drop of glue between the head, first and second antennal segments has little to no effect on several measures of olfactory navigation behavior. The probability of movement is slightly increased both during and before/after odor delivery, but this increase is not significant.

**Supplementary Figure 2 (related to Figure 2):**
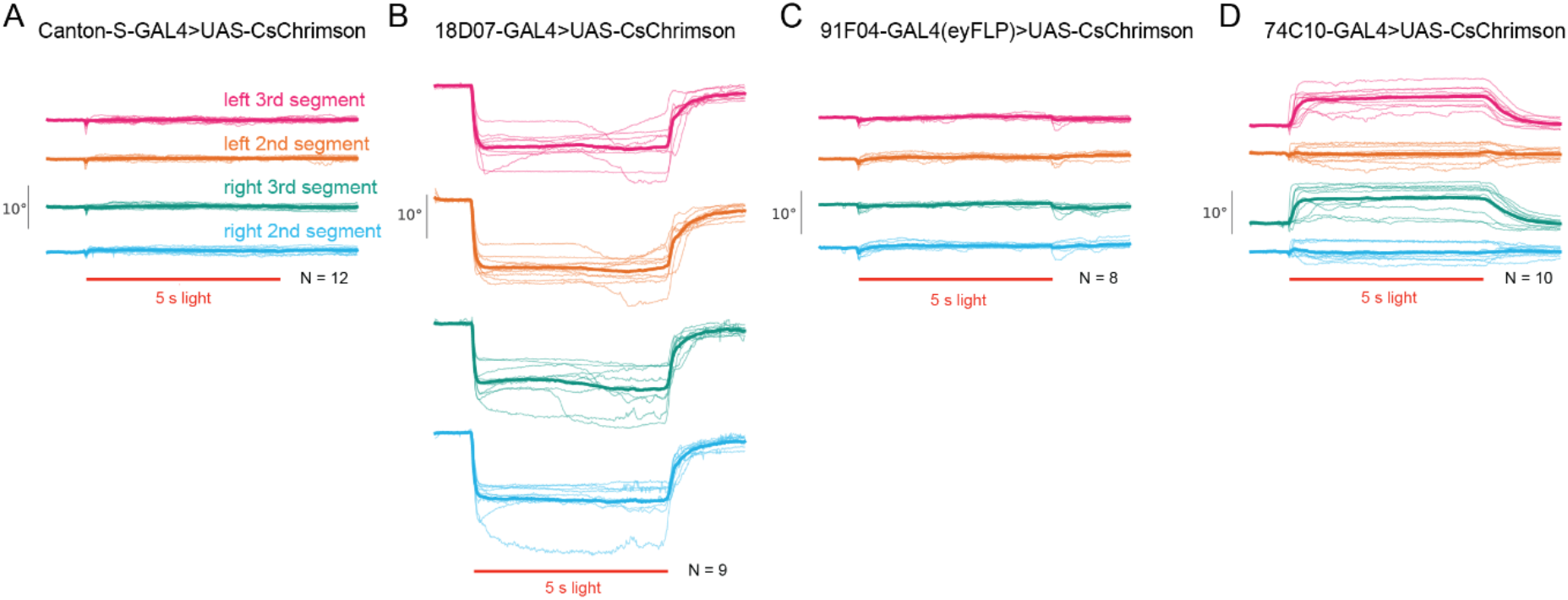
(A) Response to light activation in control flies (*Canton-S > Chrimson*, N = 12 flies) for left and right 2^nd^ and 3^rd^ antennal segments. (B) Second and third antennal responses to light activation of the first motor neuron line (*18D07 > Chrimson*, N=9 flies). (C) Second and third antennal responses to light activation of the second motor neuron line (*91F02 > Chrimson*, N=8 flies). (D) Second and third antennal responses to light activation of muscle 3 (*74C10 > Chrimson*, N=10 flies).

**Supplementary Figure 3 (related to Figure 3):**
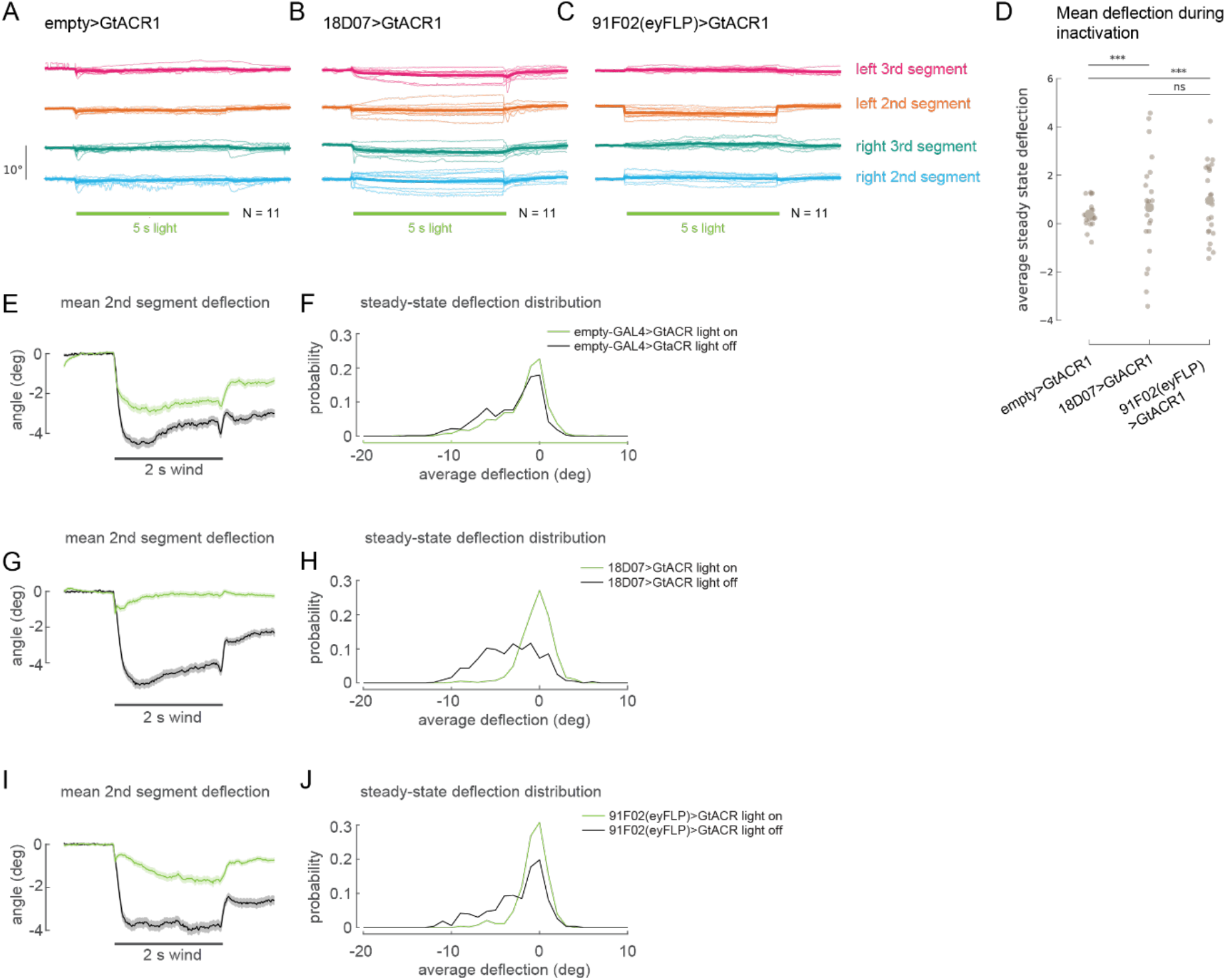
(A-C) Antennal deflections produced by light inactivation in each genotype. Green bar indicates when 5 s light stimulus was delivered. This lines represent single fly averages; thicker line indicate cross-fly averages. *Empty-Gal4>GtACR*: N= 11 flies; *18D07>GtACR*: N= 11 flies; *91F02>GtACR*: N= 11 flies. (D) Average steady-state deflection during the last two seconds of inactivation for single flies (smaller circles) and across flies (larger circle). Distribution of deflections in *18D07>GtACR1 and 91F02>GtACR1 were* different from *empt-Gal4y>GtACR1* (p=0.0001 and p<0.0001, respectively, Levene’s test) but not from one another (p=0.36). (E-J) Left: time course of average 2^nd^ segment deflections in each genotype during light (green) vs no light (black) for each genotype. Right: Distribution of steady-state antennal deflections during light (green) vs no light (black) for each genotype. Error bars represent standard error of the mean. We observed a small decrease in mean antennal displacement in control flies in response to light, but no change in deflection time course.

**Supplementary Figure 4 (related to Figure 4):**
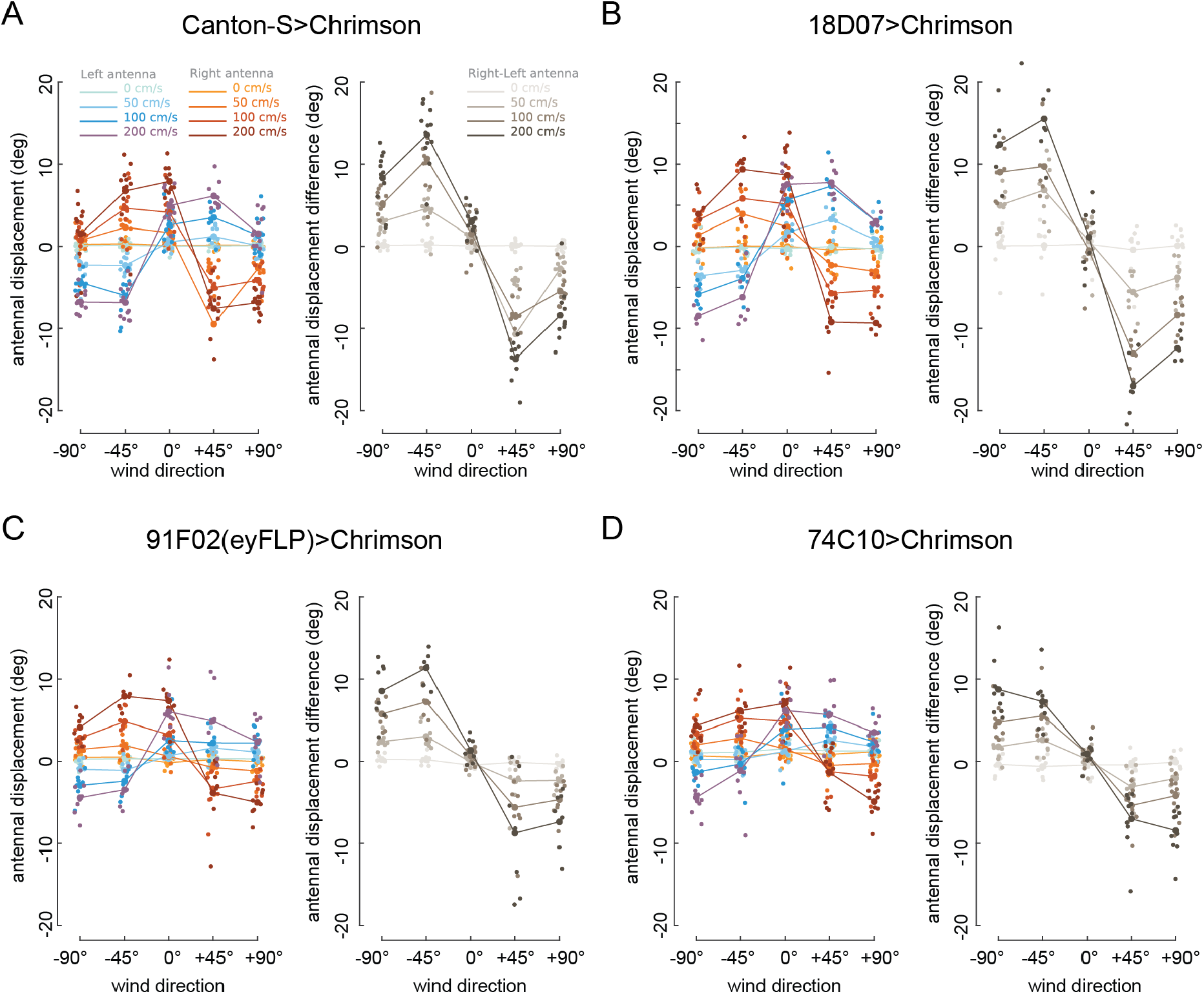
Steady-state antennal deflections for each antenna across wind directions and speeds for each genotype. Left: blue-purple hues indicate left antennal responses, orange-red hues indicate right antennal responses. Right: difference between right and left antennal deflections shown in gray hues. Smaller dots represent single flies. Lines connect represent cross-fly averages (larger dots). (A) CantonS>Chrimson, N=12 flies. (B) *18D07>Chrimson*, N=9 flies. (C) *eyFLP>GAL80;91F02>Chrimson*, N= 8 flies. (D) *74C10>Chrimson*, N= 10 flies.

## STAR Methods

### RESOUCE AVAILABILITY

#### Lead contact

Further information and requests for resources and reagents should be directed to and will be fulfilled by the lead contact, Katherine Nagel.

#### Materials availability

This study did not generate new unique reagents.

#### Data and code availability

Data generated in this study will be made available on Zenodo upon publication. Code generated during this study will be made available on Github.

## EXPERIMENTAL MODEL AND SUBJECT DETAILS

For all experiments, we used adult *Drosophila melanogaster*, between 1 and 10 days old (typically 2-4 days old). For activation experiments (Canton-S>Chrimson, 18D07>Chrimson, 91F02>Chrimson, and 74C10>Chrimson), we used female to male ratios of 7:5, 5:4, 4:4, and 8:2, respectively. We observed no obvious difference in antennal movements between male and female flies. For all other experiments, we used only female flies. Exact genotypes for each figure panel are listed in the table below. Parental strains and RRIDs are listed in the Key Resources Tables.

### Fly strains

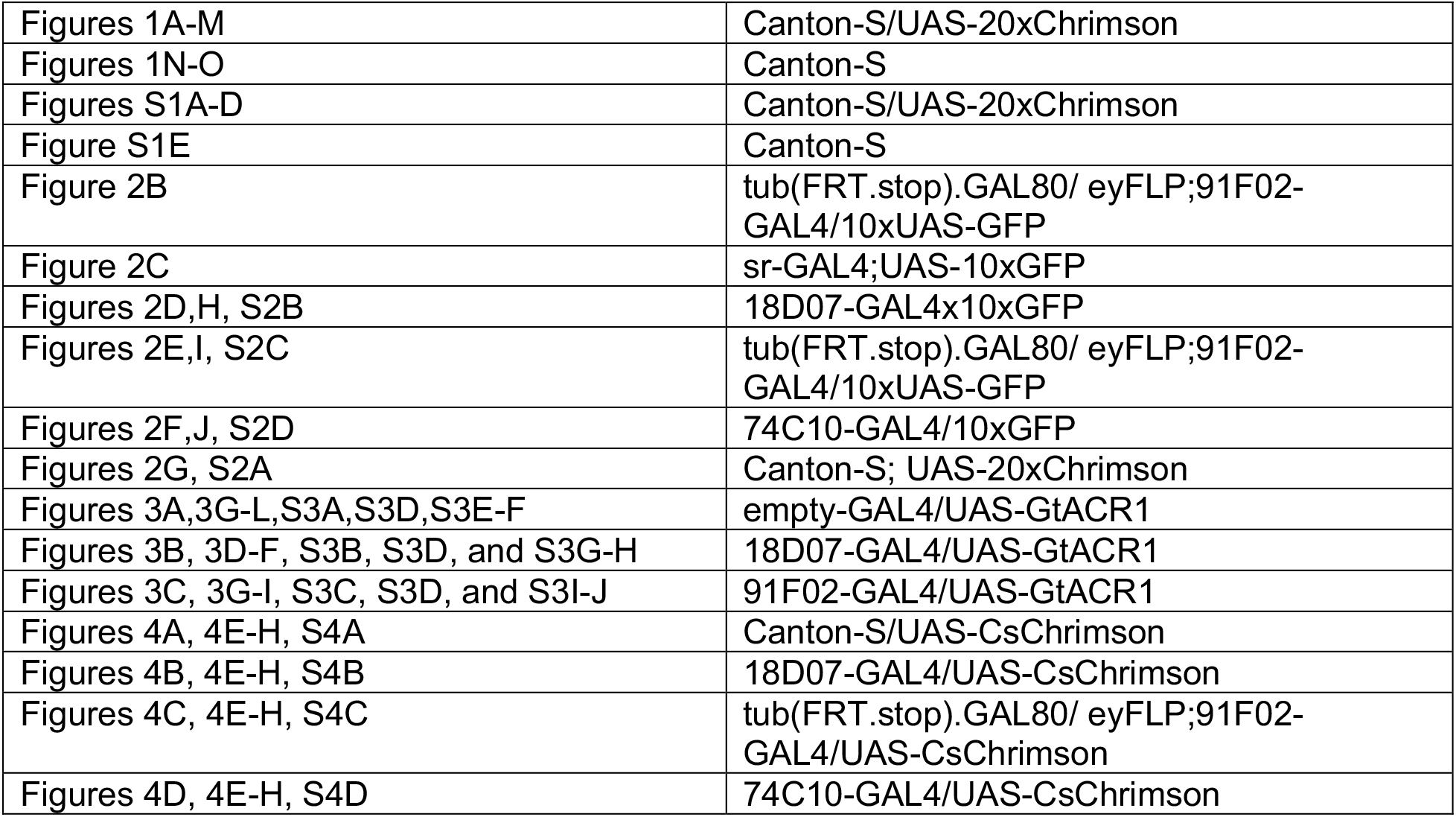

## METHOD DETAILS

### Wind Apparatus and Antenna Tracking

To mount flies in the behavior apparatus, we cold-anesthetized flies and cut off each leg at the femur-trochanter joint. Next, we glued the anterior edge of the thorax and the dorsal rim of the head to the edges of a thin stainless steel cutout attached to a plastic holder using a small amount of UV-cured glue. This holder was positioned in the center of a wind manifold with channels directed to the fly from 5 directions (−90°, -45°, 0°, +45°, and +90° relative to the fly’s midline). Wind speed was controlled using a mass flow controller (Aalborg GFC17A-VAL6-A0) and the timing and direction by a series of solenoid valves (Lee LHDA1233115H) controlled by an Arduino MEGA 2560 and custom Matlab software. We measured wind speed with a calibrated hot wire anemometer (Dantec MiniCTA with wire probe).

For odor delivery experiments, we used the same wind manifold but with an additional set of valves controlling odor delivery (same as used in a previous study: Suver et al., 2019). For these experiments, we delivered a 2 s wind pulse (60 cm/s) to 3-4 day old female flies from -45 or +45 deg. This wind contained either humidified air or 1% apple cider vinegar (Pastorelli brand).

We measured movements of the antennae with a camera mounted in front of the fly (head-on) at a framerate of 60 Hz. The fly was illuminated by IR light (880 nm) delivered by two bare fiber optic cables aimed towards the left and right of the fly from behind (Thorlabs). Wingbeats were measured using reflected IR light detected by a tachometer affixed with an IR filter (https://github.com/janelia-kicad/light_sensor_boards) mounted below the fly.

To track movements of the antennae, we used DeepLabCut ^46^ for pose estimation (version 2.0.5.1, Mathis et al., 2018). We trained our network using a set of 59 frames from 35 flies across genotypes. After an initial training of the network using 30 randomly selected frames, we selected a few additional frames by hand from videos in which we observed more tracking errors (e.g., frames when the fly was flying or those with particularly large antennal movements). We tracked 32 points on the head of the fly, including several stationary points on the head to determine the head axis, and several along the second and third segments of the antenna (Fig. S1A). We set a high training/test fraction of 0.95 (95% frames used for training, 5% for test) because our dataset was relatively unvarying (head-fixed flies with similar lighting). We used a ResNet-50-based neural network, and our network training yielded a test error of 3.48 pixels and train error of 1.31 pixels in an image size of 640×480 pixels.

### Anatomy

For the antenna image in Figure 2A, we placed a freshly removed pair of antennae from the head still attached to cuticle using fine forceps, and placed these on a damp Kimwipe. We took this image using a Google Pixel 4a aimed by hand into one eyepiece of a stereomicroscope (Leica M80).

To visualize antennal muscles and the motor neurons, we dissected whole antennae in PBS using fine forceps. We took care to avoid damaging the antennal nerve, leaving them connected to central brain. After removal, we placed the antennae in 4% paraformaldehyde solution (in PBS) for 50 minutes on an oscillator at room temperature. Following this, we washed the antennae in PBST for 15 minutes, followed by a wash in PBS for another 15 minutes. To stain antennal muscles, we placed antennae in a solution of PBS, Alexa Fluor Phalloidin 568 (10µL stock in 1 mL PBS, Invitrogen A12380) and rabbit anti-GFP 488 (10µL in 1mL PBS, Thermofisher A6455) for approximately 48 hours. We then removed the antennae from the solution and placed it in PBS for 15 minutes on an oscillator at room temperature. Afterwards, we mounted the antennae anterior side up on a glass microscope slide in Vectashield (Vector Labs H-1000-10), covered with one #0, 22 × 22 mm coverslip, and sealed with nail polish. We imaged antennae at 20x magnification on a Zeiss LSM 800 confocal microscope at 1 µM depth resolution. Final images are presented at maximum z-projections over relevant depths.

### Optogenetics

We placed adult flies on food enriched with all-trans retinal (50 µL all-trans retinal in 1 tsp rehydrated potato flakes placed on top of standard cornmeal-molasses fly food) for 24-48 hours prior to optogenetics experiments. For Chrimson activation, we used red light with an intensity sufficient to produce antennal deflections in our initial motor neuron screen (3.23 µW/mm^2^ measured at 658 nm at the location of the fly when mounted). For GtACR1 inactivation experiments, we used green light at 70 µW/mm^2^ measured at 530 nm.

### Walking arena behavior

For walking behavior, we used wild type Canton-S flies that were 2-5 days old (roughly equal numbers of males and females). These flies were raised on standard food and a 12:12 light cycle at 25°C. These flies were then starved (housed in an empty vial with a damp kimwipe) for 24 hours prior to the experiment to encourage odor-seeking. We blocked movements of the first and second antennal segments by applying UV-curing glue to a tiny portion of the head to the first segment and dorsal edge of the second segment in approximately rest position. For each fly, after the experiment finished, we determined whether our gluing procedure was successful by verifying under a light microscope that the third segment was free but first and second remained fixed to the head. We used previously described walking wind tunnels^50^ to present controlled wind and odor stimuli to freely walking flies. We placed cold-anesthetized single flies in individual walking tunnels. We let these flies acclimate for several minutes, then presented them with either no wind, wind, or constant wind with a 10 s pulse of 10% apple cider vinegar (made with Pastorelli brand).

### QUANTIFICATION AND STATISTICAL ANALYSIS

We analyzed all data in Python 3.7.6 and Matlab 2018b.

To analyze active antennal movements, we first computed 2^nd^ and 3^rd^ segment angles with respect to the midline based on the points shown in Fig 1B. Transient tracking errors (angles greater than 35 deg, which we only ever observed as being caused by tracking errors) were detected and omitted from analysis.

Mean second segment displacements were computed by averaging across all trials for a given stimulus condition. In Figures 3D and 3E, we averaged across all directions at a windspeed of 200 cm/s. Distributions of steady-state second segment displacements were computed by taking the average antennal deflection in the difference between baseline (average 0.75 s before light on) and steady-state light response (average of the last 1.25 s during light stimulation).

To detect rapid antennal movements, we first baseline subtracted each angular measurement (average angle during the first second of the trial during which there was no light or stimulus) and low-pass filtered with a butterworth filter (cutoff frequency = 10 Hz). We then found peaks of in the third segment angle trace using signal.find_peaks with prominence of 1.5 degrees. We omitted transient movements at the light or wind stimulus on and offset (within 2 frames at onset for both, and 8 or 10 frames at offset, respectively). We used third segments in this analysis because active movements were visible in both second and third segment traces and typically had larger signal to noise in the third segment trace (see Fig. S1B, C). We detected a small number of rapid active movements from second segment traces and detected no statistically significant difference in counts of active movements obtained from third vs second segment in this sample (Fig. S1C).

For active movement peri-stimulus time histograms (Figures 1J, 3I, and 3L), we mirrored left antennal movements and pooled these with data from the right antenna. We normalized histograms to the number of trials, converted to movements per s, and low-pass filtered with a Butterworth filter with cutoff frequency of 5 Hz. We computed auto- and cross-correlograms across all raw antennal movements for the left or right second or third segment, relatively, using scipy.signal.correlate. To compare the variation in this data set, we computed the mean cross-correlation for each fly, and compared across conditions with a paired student’s t-test.

Most statistical analyses were performed using a paired two-sided student’s t-test (scipy.stats.ttest_rel). For Chrimson activation quantification (Figure 4F), we used an unpaired two-sided student’s t-test (scipy.stats.ttest_ind) test as we were comparing across genotypes. To compare the distribution of antennal deflections during inactivation, we used Levene’s test to assess variance (scipy.stats.levene; Fig. S3D).

To compute the Fisher Information, we first fit a smooth tuning curve, d_smooth(φ) to the plot of mean antennal displacement difference (in degrees) as a function of wind direction for each genotype (also in degrees, using 200 cm/s wind) with one fly left out for jackknifing (see below). The tuning curve had the form of a Gaussian function multiplied by a ramp:

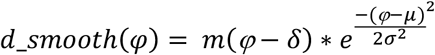

where *φ* is the wind direction, and *m, δ, μ*, and *σ* are free parameters. The parameters were determined by minimizing the mean squared error between this tuning curve and the experimentally measured values of d(φ). The Fisher Information was computed as the mean across flies of

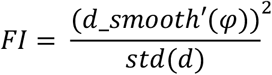

The numerator was computed by differentiating d_smooth and squaring. The denominator was computed by first computing the variance of d for each wind direction, then averaging across wind directions and flies to obtain a single measurement of variance. The standard deviation was the square root of that quantity. The mean and error bars of FI, the quantities plotted in Figure 4H, were computed by jackknifing across flies, using the formula:

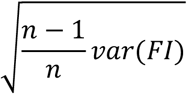

where *n* is the number of flies.

We measured movement parameters for walking behavior using custom Matlab scripts as previously reported ^30,50^. Trials in which flies moved less than 25mm total or for less than 5 trials were omitted from our analysis. Groundspeed was computed by taking the distance between adjacent samples divided by the 20 ms frame interval. Probability of moving was the probability of groundspeed exceeding 1 mm/s. Upwind velocity was the absolute value of the difference in y-coordinates divided by the frame interval. Angular velocity was computed by dividing the absolute value of the difference in unwrapped orientation by the frame interval. Curvature is plotted as the angular velocity divided by the groundspeed. Variance is presented as standard error of the mean.

## KEY RESOURCES TABLE

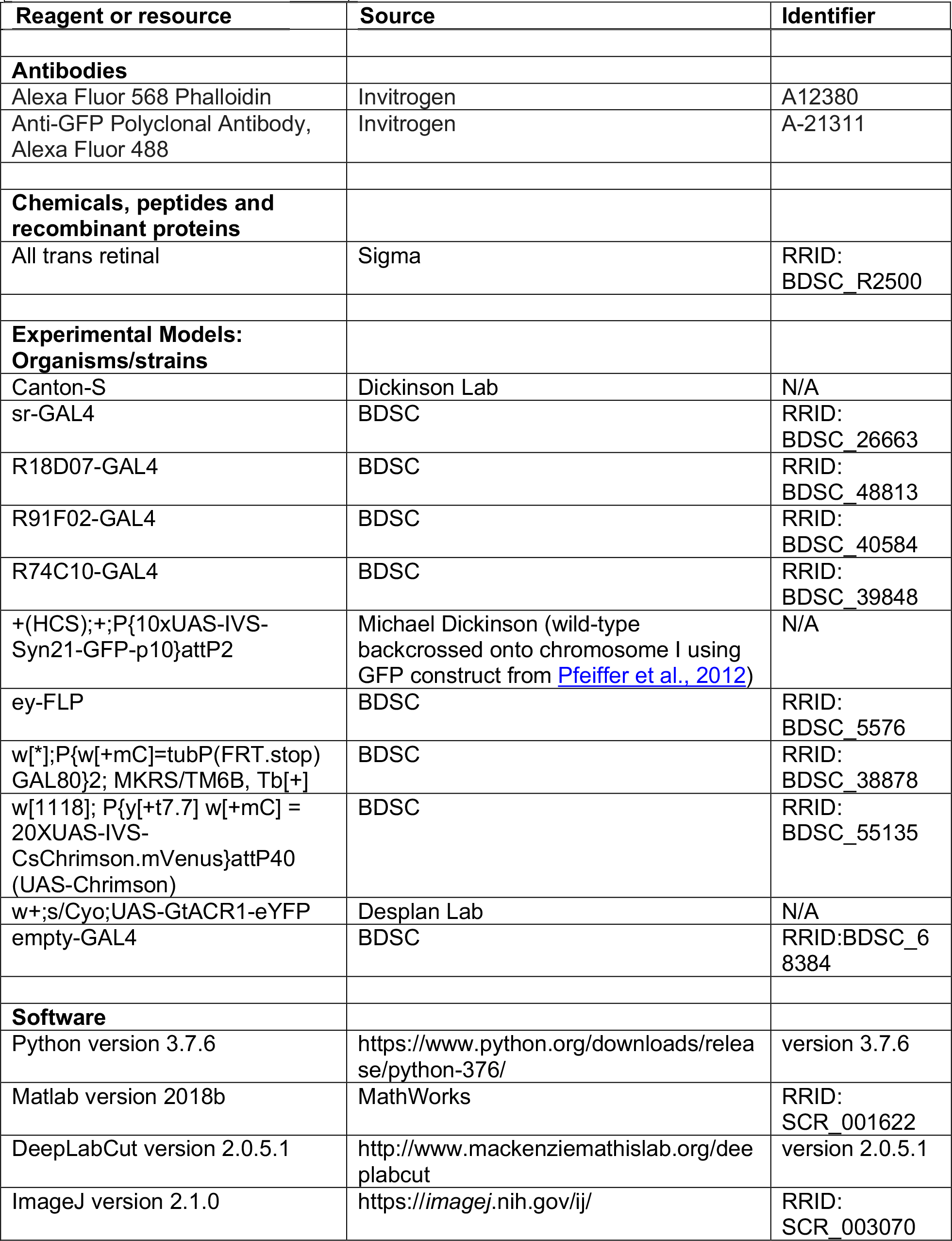

